# Percolate: an exponential family JIVE model to design DNA-based predictors of drug response

**DOI:** 10.1101/2022.09.11.507473

**Authors:** Soufiane M.C. Mourragui, Marco Loog, Mirrelijn van Nee, Mark A van de Wiel, Marcel J.T. Reinders, Lodewyk F.A. Wessels

## Abstract

**Motivation:** Anti-cancer drugs may elicit resistance or sensitivity through mechanisms which involve several genomic layers. Nevertheless, we have demonstrated that gene expression contains most of the predictive capacity compared to the remaining omic data types. Unfortunately, this comes at a price: gene expression biomarkers are often hard to interpret and show poor robustness.

**Results:** To capture the best of both worlds, i.e. the accuracy of gene expression and the robustness of other genomic levels, such as mutations, copy-number or methylation, we developed Percolate, a computational approach which extracts the joint signal between gene expression and the other omic data types. We developed an out-of-sample extension of Percolate which allows predictions on unseen samples without the necessity to recompute the joint signal on all data. We employed Percolate to extract the joint signal between gene expression and either mutations, copy-number or methylation, and used the out-of sample extension to perform response prediction on unseen samples. We showed that the joint signal recapitulates, and sometimes exceeds, the predictive performance achieved with each data type individually. Importantly, molecular signatures created by Percolate do not require gene expression to be evaluated, rendering them suitable to clinical applications where only one data type is available.

**Availability:** Percolate is available as a Python 3.7 package and the scripts to reproduce the results are available here.

## 1 Introduction

Over the course of their lifespan, human cells accumulate molecular alterations that result in the modification of cell behavior [27]. When aggregated at the tissue level, these alterations can compromise tissue homeostasis, in turn clinically impacting a patient [13]. Understanding the combined effect of these alterations is key to designing bespoke lines of treatment [28, 33]. These molecular alterations occur at different genomic levels and are recorded using different technologies, collectively referred to as “omics” technologies. Each of these omic measurements offers only partial information regarding the compromised tissue. Aggregating different omic measurements, an analysis known as multi-omics integration, is therefore necessary to generate a comprehensive picture of the molecular features underlying a cancerous lesion [5, 20].

Owing to their high versatility, cell lines offer a cost-effective model system for drug response modelling [8]. Specifically, large scale consortia have industriously subjected a large number of cell lines to hundreds of different compounds, yielding valuable drug response measurements [16, 12, 32]. A key challenge resides in combining these response measurements with multi-omics data to study mechanisms of resistance and sensitivity[24]. Existing approaches focus on combining all omics data types and can be ordered based on the stage of the analysis at which the integration is performed [6]. At one extreme, early integration approaches [4, 19] first aggregate all features from all data types to process them all simultaneously. At the other extreme, late integration approaches first compute a representation of each data type individually, and subsequently combine these representations [11, 26, 36]. Several other methods can be positioned along this ordering, and differ by the analysis stage during which the grouping of data types is performed [41]. Although promising and encouraging, these methods do not take into account the quality of the data types and do not explicitly model their topology [2], i.e., how the data types relate to each other regarding information content and capacity to predict drug response. In particular, it has been observed that, although it has traditionally been the least clinically actionable data type, gene expression consistently prevails over other data types [9] and provides similar performance as early-integration approaches [1], obviating the need for complex integration strategies.

In order to maintain the predictive power of gene expression data, while exploiting the robustness of the most actionable data types, we present Percolate, an unsupervised multi-omics integration framework. Percolate sets itself apart from other integration approaches as it aims to eliminate gene expression measurements from the final predictor, rather than integrating it with all other data types. This is achieved by extracting the joint signal between gene expression and the other data types in an iterative fashion. First the joint signal between gene expression and Data type 1 (e.g. mutations) is extracted. Then the remaining signal (not shared with Data type 1) is employed to extract the joint signal of gene expression with Data type 2 (e.g. copy number data). This procedure is repeated for all omics data types. In this way, the gene expression signal is “percolated” down the other omics data types, ideally extracting all predictive signal from the gene expression data. Technically, Percolate employs a popular framework, called JIVE [11, 26], which breaks down paired datasets into joint and individual signals. We first extended JIVE to non-Gaussian noise models employing GLM-PCA [7]. Specifically, we used an alternative optimization, the decomposition of saturated parameters [21], which we theoretically proved to be competitive with the original formulation. Finally, we developed an out-of-sample extension for JIVE, useful when only one of the two data types is available.

We first show that comparing gene expression to other data types individually recovers a known topology of multi-omics data. We then show that the information shared between individual omic data types and gene expression increases drug response predictive performance for the individual omic data types. Finally, reconstructing the joint signal solely from mutation, copy-number and methylation, we show that the signatures derived from “percolating” gene expression down these data types recapitulate the drug response predictive performance of these data types.

## 2 Methods

### 2.1 Trade-off between robust and predictive types

We consider four data types: mutations (MUT), copy number aberrations (CNA), methylation (METH) and gene expression (GE). MUT and CNA directly measure genetic aberrations and therefore rely on DNA measurements. Due to several biological and technological factors, these measurements are highly robust and suffer from little technical artefacts. On the other end of the spectrum, GE measures RNA abundance, a process known for exhibiting large biological variability and prone to technical artefacts.

Between these two extremes, methylation offers an intermediate level of robustness. However, when it comes to drug response prediction, the order is reversed: GE offers, on average, a better predictive performance than METH, and significantly outperforms MUT and CNA [1, 8, 17]. This leads to a trade-off between robustness and predictive ability (Figure 1A) with MUT and CNA being the most robust and least predictive and GE being the most predictive and least robust, with METH rating at the intermediate level in terms of robustness and predictive capacity.

**Figure 1:**
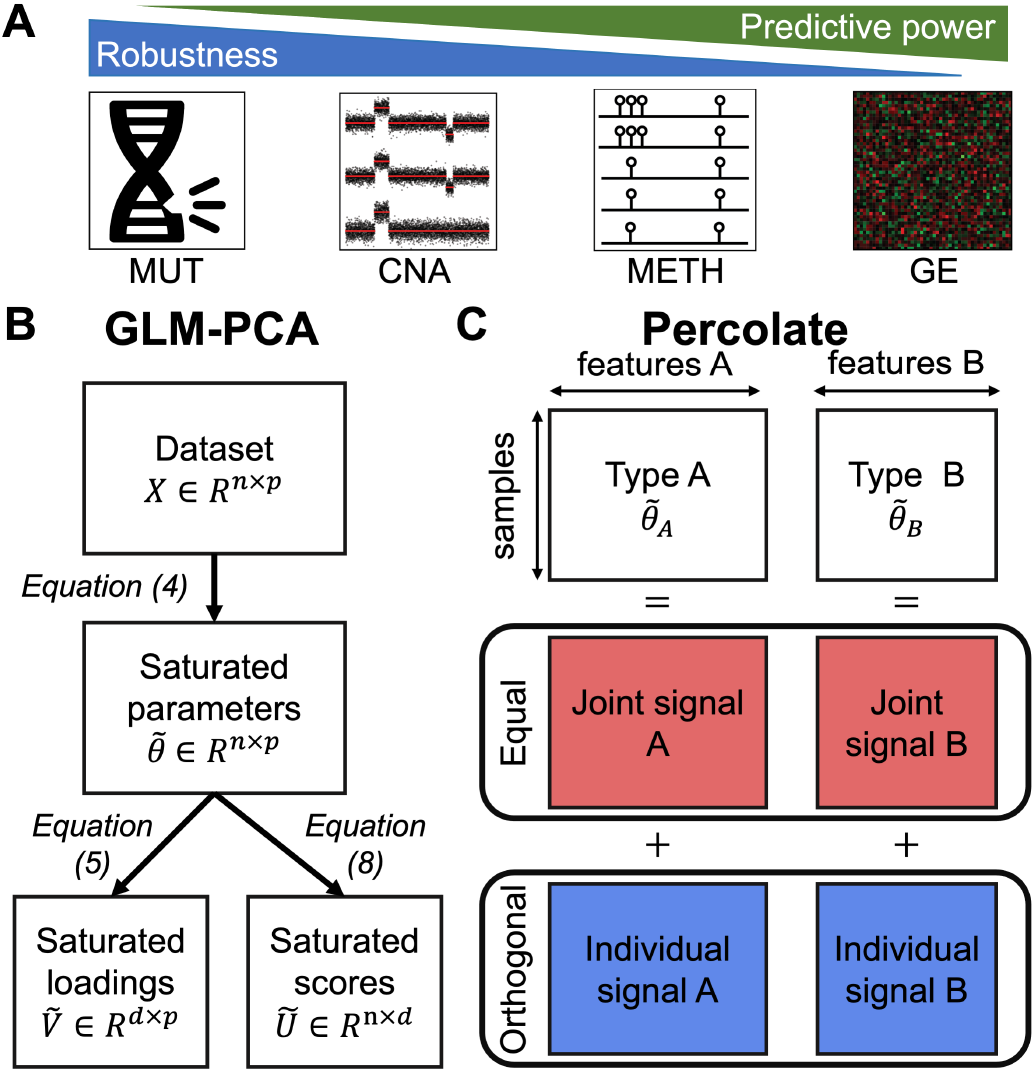
Dissecting multi-omics topology using Percolate bridges the gap between predictive and robust data types. (**A**) Trade-off between robust data types (MUT, CNA) and predictive types (METH, GE). (**B**) Workflow of our implementation of GLM-PCA, which relies on the projection of saturated parameters. (**C**) Workflow of Percolate, which extends JIVE to non-Gaussian settings by comparing the low-rank structures of saturated parameter matrices.

### 2.2 Exponential family distribution

Our integrated approach is inspired by AJIVE [11], a computational approach which takes as input two paired datasets and computes a joint and a data-specific signals. AJIVE is an extension of the JIVE model [26], which we selected, among other extensions [35, 37], for its computational tractability and its mathematical formulation which is amenable to the derivation we propose. JIVE, AJIVE, and derivations thereof, critically rely on Principal Component Analysis (PCA) which assumes a Gaussian noise model on the data [22, 39]. To extend this framework to non-Gaussian settings, we make use of a generalized formulation that can deal with a wider class of parametric distribution models, i.e., the so-called exponential family [31].

#### Definition 2.1

(Exponential family distribution). *Let* 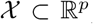, *we say that a random vector* 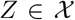 *follows an **exponential family distribution** if its probability density function f can be written as*

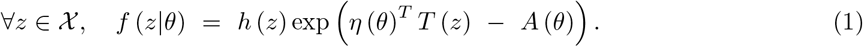

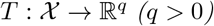 is called the **sufficient statistics**, *θ* ∈ ℝ^*q*^ *the **exponential parameter**, η*: ℝ^*q*^ →∈ ℝ^*q*^ *the **natural parametrization**, A*: ℝ^*q*^ → ℝ *the **log-partition function** and h*: 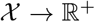 *the base measure*.

The exponential family encompasses a broad set of distributions (Supp. Table 1), including the Gaussian distribution with unit variance, the Poisson, the Bernoulli, the Beta or the Gamma distributions. Practically, the functions *A, T* and *η* are modelling choices which can be tuned for any specific application.

### 2.3 Saturated model parameters

For this section, we consider a data matrix *X* ∈ ℝ^*n×p*^, with *n* (resp. *p*) the number of samples (resp. features). We model this data using an exponential family distribution 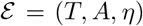 (Definition 2.1), which choice is motivated by prior knowledge. For instance, if the data is known to be binary, one would turn to 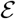 defined by the Bernoulli distribution, while another data distribution would lead to a different choice of functions (Supp. Table 1). We denote by *q* the dimensionality of *T* and *A* output space.

#### Definition 2.2

(Negative log-likelihood). *We define the **negative log-likelihood**, denoted 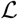, as follows*:

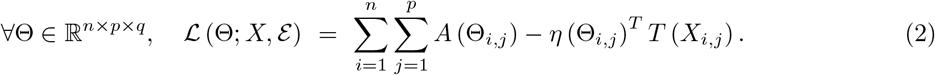

#### Definition 2.3

(Saturated parameters). *We define the saturated parameters* 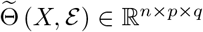 *as the minimizers of 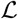, i.e*.,

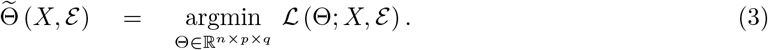

The saturated parameters correspond to single-sample maximum likelihood estimates. This quantity, which will be the pillar of our approach to GLM-PCA (Section 2.4), can be computed as follows.

#### Proposition 2.4

(Computation of saturated parameters). *Assume that A and v are differentiable with invertible differentials. Then, denoting J as the Jacobian of a function*:

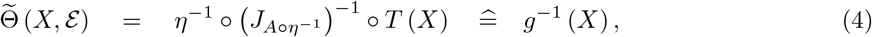

*using an element-wise operation on all elements of X*.

*Proof*. We refer the reader to the Supplementary Material (Section 4) for the proof.

Proposition 2.4 shows that the saturated parameters correspond to a dual representation of the data motivated by prior knowledge on the data-distribution. We will exploit this representation *à la PCA* to find the main sources of variations in a framework called GLM-PCA.

### 2.4 Generalized Linear Model Principal Component Analysis (GLM-PCA)

JIVE is based on Principal Component Analysis (PCA), which admits three equivalent definitions: maximization of projected variance, minimization of reconstruction error and maximization of a Gaussian likelihood with unit-variance. This latter definition can be restrictive for non-Gaussian data and we therefore set out to replace PCA by an extension called **GLM-PCA** [7]. In these methods, the Gaussian likelihood is replaced by an exponential family distribution. The original approach from *Collins et al* [7] minimizes a negative log-likelihood using an SVD-like decomposition for the exponential parameters, yielding three different matrices. Refinements of this idea, which solve a similar optimization problem, have been proposed in the literature [23, 25] and offer competitive routines for the computation of these three matrices. Another take on this problem, which relies on the projection of saturated parameters, has recently been developed by *Landgraf et al* [21]. This approach offers the advantage of a simpler single-matrix optimization instead of concomitantly optimizing on three. Furthermore, the out-of-sample extension relies on a matrix multiplication and is thus computationally fast. These two approaches therefore aim at finding the same decomposition through different computational routines. We here present these two approaches and prove that the latter offers a similar or better minimizer for the negative log-likelihood, which, to the best of our knowledge, had not been established.

#### 2.4.1 Two formulations of GLM-PCA

##### Definition 2.5

(SVD-type [7]). *SVD-type GLM-PCA computes three matrices*, 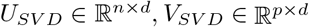 and 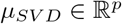 (*diagonal), alongside a vector* 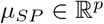 *defined as*

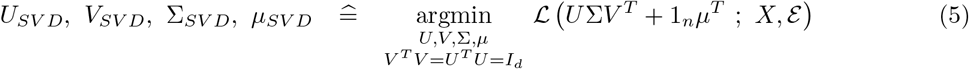

##### Definition 2.6

(Projection of saturated parameters [21]). *GLM-PCA by projection of saturated parameters computes one matrix, V_SP_* ∈ ℝ^*p×d*^ *alongside a vector μ_SP_* ∈ ℝ^*p*^ *defined as*

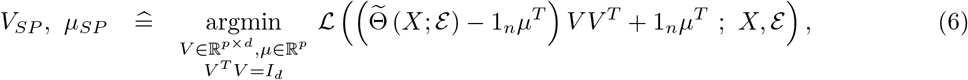

The loading matrices (*V_SVD_* and *V_SP_*) and the score matrix (*U_SVD_*) have orthogonal constraints, which is similar to PCA where scores are by construction uncorrelated.

#### 2.4.2 Equivalence of the formulations

We here show that the projection of saturated parameters provides a competitive minimization when compared to the SVD-type decomposition. The main result is based on Supp. Lemma 5.1 and we refer the reader to the Supplementary Material (Section 5) for a complete proof.

##### Theorem 2.7.

*Let us define *U_SVD_,V_SVD_, Σ_SVD_* and *μ_SVD_* as in Definition 2.5, and *V_SP_, μ_SP_* as in Definition 2.6. The likelihood resulting from the two optimization processes satisfies*

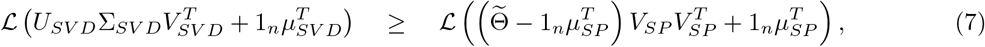

*where the dependencies on X and* 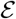 for 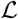 *and* 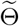 *were removed for verbosity’s sake*.

Theorem 2.7 shows that, although the two approaches compute the same decomposition, the one obtained from saturated parameters yields a lower or equal negative log-likelihood. It is also worth noting that the SVD-like optimization is usually performed by alternate optimization [40] and the initialization can play a major role in the convergence. The projection of saturated parameters only requires one minimization round, and is thus faster and less prone to initialization effects. Using the decomposition of saturated parameters, however, comes at a price: there is an infinity of solutions, all equal up to a unitary transformation. In order to obtain sample scores that are uncorrelated, we proceed as follows.

##### Definition 2.8

(Sample scores). *Let V_SP_ and μ_SP_ be defined as in Definition 2.6 and assume that rank* 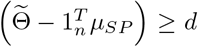. *Then rank* 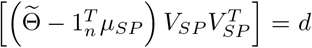 *and we define U_SP_*, Σ_*SP*_ *and W_SP_ as the unique rank-d SVD decomposition of the saturated parameters, i.e*.

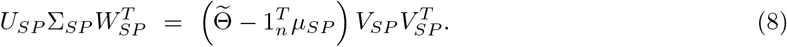

It is worth noting that the equality in Equation 8 is not an approximation and this second SVD does not entail any loss of information. It is a pure computational maneuver to whiten the obtained scores.

#### 2.4.3 Hyper-parameter optimization

The solution of Equation (6) is an optimization problem with a Stiefel-manifold constraint, which we solved by using recent advances in auto-differentiation [30] and optimization on Riemmannian manifolds [29]. We modelled the functions *A, T* and the negative log-likelihood using PyTorch; stochastic gradient descent (SGD) on the Stiefeld-manifold was performed using McTorch. Such a formulation allows to employ a large variety of exponential family distributions without the need for heavy and potentially cumbersome Lagrangian computations. Our optimization scheme relies on four hyper-parameters: number of factors (or principal components), learning rate, number of epochs and batch size. To determine them, we compute the Akaike Information Criterion (AIC) of the complete data for various values of d and different hyper-parameters [3]. For a GLM-PCA model with d PCs, the AIC corresponds to the sum of the data log-likelihood and the number of model parameters, which we estimate as the dimensionality of the Stiefel manifold 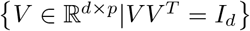, equal to *p_d_* – *d*(*d* + 1)/2. Among all trained models, we select the one which harbors the smallest AIC.

### 2.5 Comparison of GLM-PCA directions by Percolate

**Setting**: We consider two datasets 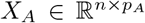 and 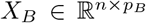 with paired samples (rows) but potentially different features. We first perform GLM-PCA independently on *X_A_* and *X_B_* using two different exponential family distributions, yielding *d_A_* and *d_B_* factors, respectively denoted as 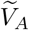 and 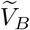. We furthermore denote by 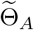 and 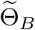 the saturated parameters of datasets A and B respectively, and 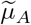 and 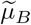 the intercept terms. Using the decomposition presented in Definition 2.8, we furthermore define 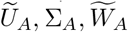 and 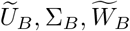.

#### Definition 2.9.

*To compare the two sets of samples scores, 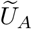 and 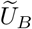, we aggregate them in a matrix* **M**, *which we decompose by SVD*:

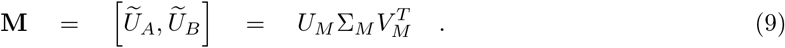

The top left-singular vectors correspond to sample scores which are highly correlated between 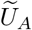 and 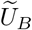, since both of these two matrices are consisting, by construction, of uncorrelated factors. Following the same intuition as in AJIVE, these can be understood as the **joint signal**, motivating the following definition.

#### Definition 2.10

(Joint and individual signals). *Let r_J_* < min (*d_A_,d_B_*), *we define the **joint signal** as the matrix* 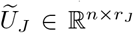 *with the top r_J_ left-singular values of* **M**. *We furthermore denote by* Σ_*J*_ *the diagonal matrix with the top r_j_ singular values of* **M**.

*We define the **individual signal** of A (resp. B), denoted as* 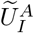 (*resp*. 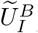), *as the signal from* 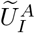 (*resp*. 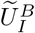) *not present in* 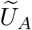 (resp. 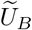), *formally*:

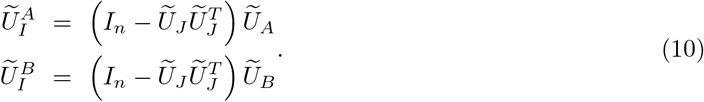

We call the complete process **Percolate**, and a summarised workflow can be found in Figure 1B-C.

In order to set the number of joint components *r_J_*, we employ a sample-level permutation scheme. We first independently permute the rows of 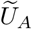 and 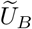, which we then aggregate as in Equation (9) to obtain the singular values. We perform 100 such permutations independently and retrieve the first singular value for each. Finally, we set *r_J_* as the number of elements in Σ_*M*_ over one standard deviation from the mean of the permuted singular values (Figure 2A).

**Figure 2:**
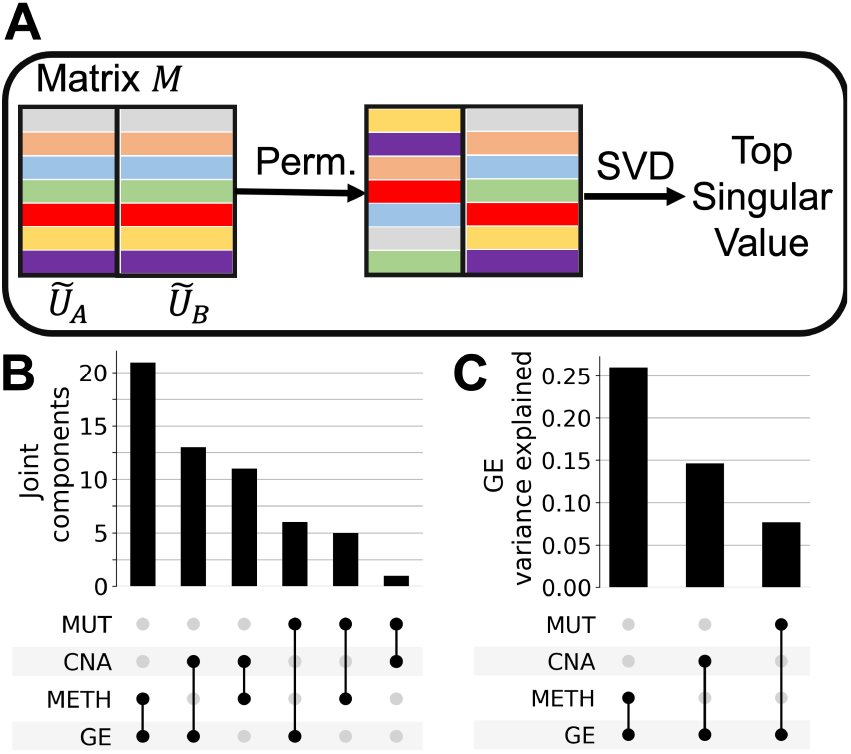
Assessing the number of joint components. (**A**) Schematic of the samplelevel permutations we perform to estimate the number of joint components. (**B**) Venn-diagram of the number of joint components obtained using the permutation scheme. (**C**) Ratio of variance explained for the GE saturated parameters matrix after projection on the joint components.

### 2.6 Projector of joint signal

AJIVE does not provide an out-of-sample extension, and we here propose a derivation thereof by rewriting the matrix *U_J_* as a function of the saturated parameters.

#### Theorem 2.11.

*Let’s decompose the matrix V_M_ as* 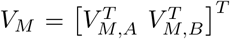 *such that* 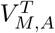 *contains the first d_A_ columns of* 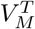 *and* 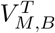 *the last d_B_ ones, we obtain*:

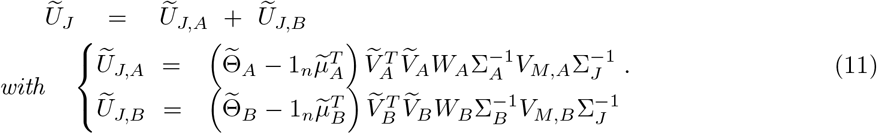

*Proof*. We refer the reader to the Supplementary Material (Section 6) for the complete proof.

The formulation of 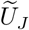 presented in Equation (11) highlights the additive contribution of both dataset to the joint signal. At test time, both views are therefore required to estimate the joint signal. To tackle the issue of missing data-view, we propose a nearest-neighbor imputation of the unknown joint-term. Let’s consider, without loss of generality, that only the view *A* is available. The joint signal has been computed using the two data matrices *X_A_* and *X_B_*, yielding 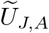 and 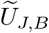. The second term contains *r_J_* terms, and we train *r_J_* corresponding k-Nearest-Neighbors (kNN) regressors. The test dataset *Y_A_* ∈ ℝ^*m×p_A_*^ can be projected on the joint signal by replacing the saturated parameter 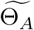 in Equation 11 with the saturated parameter of the test data. We then estimate the second term by means of the *r_J_* kNN regression models. Adding these two terms yields an estimate of the joint signal.

### 2.7 Drug response prediction

We assess the predictive performance of a dataset by employing ElasticNet [42], which has been shown, inspite of its relative simplicity, to outperform more complex non-linear models when it comes to drug response prediction [8, 17, 38]. For a given dataset, we perform nested cross-validation as follows. First, datasets are stratified into 10 groups of equal size. For each group (10%), we employ a 3-fold crossvalidation grid search on the remaining 90% to determine the optimal ElasticNet hyper-parameters (*ℓ*-ratio and penalization). We then fit this optimal ElasticNet model on the 90% to predict the class labels on the 10%. Repeating this procedure, we obtain one cross-validated estimate per sample and we define the **predictive performance** as the Pearson correlation between these estimates and the actual values.

### 2.8 Data download, modelling and processing

We consider four data types in our analysis (Table 1) which we modelled using different exponential family distributions (Supp. Material). The GDSC data was accessed on January 2020 from CellModel Passport [16]. For GE, MUT and CNA, we restricted to protein coding genes known to be frequently mutated in cancer, referred to as the **mini-cancer genome** [15]. GE was corrected for library size using TMM normalization [34] and mutations were restricted to non-silent.

**Table 1:** Exponential family distributions. Gaussian distribution is assumed to have unit variance. The dispersion parameter *r* is fixed for the Negative Binomial.

| Data type | Family distribution |
| --- | --- |
| Copy-number aberration (CNA) | Log-Normal or Gamma |
| Gene expression (GE) | Negative-Binomial |
| Methylation (METH) | Beta |
| Mutation (MUT) | Bernoulli |

## 3 Results

### 3.1 The breakdown of the joint signals highlights the topology of multi-omics data

To compare data types, we employ Percolate using the distributions defined in Table 1, and a number of PCs set using the procedure presented in Subsection 2.4 (Supp. Figure 2). For each comparison, setting the number of joint components is a crucial step, as it defines the threshold between the joint and individual signals. For that purpose, we used a sample level permutation test (Figure 2A, Subsection 2.5).

We observe that GE shares 21 joint components with METH, 13 with CNA and only 6 with MUT, which is coherent with the gradient put forward in Figure 1. We furthermore observe that MUT is consistently the data type with the least number of joint components (Figure 2B), highlighting the weakness of the signal coming from MUT data, corroborating previous measured topologies of multi-omics data [2]. To measure the strength of the underlying joint signals, we computed the proportion of GE variance explained by the joint directions (Figure 2C), computed as the ratio between the joint signal variance and the variance of the GE’s saturated parameters matrix. We observe that the joint signal between GE and METH explains 26% of GE variance, while this figures drops to 14% and 7% for CNA and MUT, respectively. These observations highlight the existence of a joint signal, of which the predictive performance can be interrogated.

### 3.2 Robust signal predictive of drug response is concentrated in the joint part

We then investigated the relevance of the joint and individual signals when it comes to drug response prediction. Considering one robust data type at a time (MUT, CNA or METH), we first decomposed the original robust data type into a signal joint with GE and an individual signal specific to the robust data type. We then computed, for 195 drugs (Methods), the predictive performance for these two signals and compared it to the original robust robust data (Figure 3A, Subsection 2.7). To ensure a proper comparison between joint, individual and cell-view, the cross-validation was performed using the same folds for all datasets.

**Figure 3.**
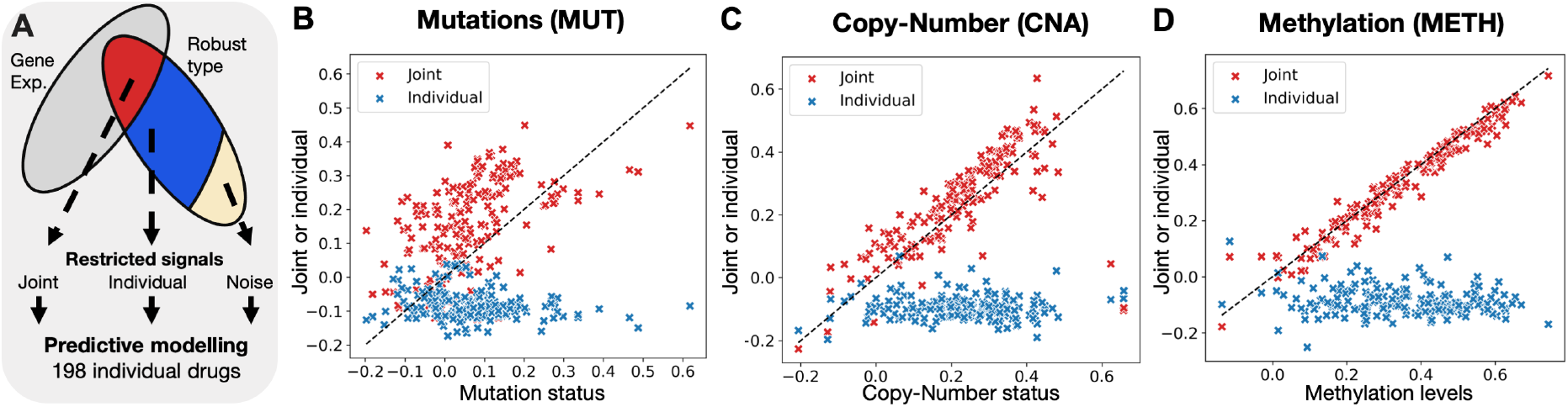
The joint signal between robust and gene expression contains most of the predictive signal. (**A**) Workflow of our approach. (**B**) Predictive performance for MUT when using Percolate between MUT and GE. Each point corresponds to a single drug, with the x-axis corresponding to the predictive performance obtained using the original mutation data, and the y-axis by either the joint (red) or the individual (blue) signals. (**C**) Predictive performance for CNA, similarly displayed as in **B**. (**D**) Predictive performance for METH, similarly displayed as in **B**.

We first analyzed the results obtained between MUT and GE data (Figure 3B). We observe that for most drugs, the predictive performance of the joint signal exceeds the predictive performance of the original robust signal, except for a number of drugs of which the response is quite well predicted based on MUT only. This set includes the drugs Nutlin-3, Dabrafenib, and PLX-4720. In contrast, the individual signal shows no predictive performance (Pearson correlation below 0) for most drugs, indicating an absence of drug response related signal in the individual portion. We then turned to CNA where the choice of distribution was unclear, with, to the best of our knowledge, no clear precedent on how to model such data. Due to the observed behavior of CNA data, we opted for two possible distributions: Lognormal and Gamma distributions (Supp. Table 1). We observe that the joint signal computed using a Gamma-distribution yields better performances than the log-normal model (Supp. Figure 3A-B). When using a Gamma distribution, a conclusion similar to the MUT data can be reached with the majority of drugs predicted well with the joint signal except three drug, AZD4547, PD173074 and Savolitinib (Figure 3C). This advocates for using the Gamma distribution for analyzing CNA data and shows that the joint signal presents an increased performance while the individual signal is not predictive. Finally, we studied the drug response performance obtained after decomposing METH using GE (Figure 3D). We observe that the joint signal presents a similar predictive performance as the original methylation data. The individual signal is, again, not predictive. These results highlight the potential of restricting predictors to the joint signal for robust data types.

### 3.3 Out-of-sample extension recapitulates the predictive performance of robust signal

In order to compute the joint signal between one robust data type and GE, one needs to have access to both modalities. However, the purpose is to become independent of non-robust GE measurements. In order to study whether the joint signal could be estimated without access to gene expression, when the predictor is applied to a test case, we exploited our out-of-sample extension (Subsection 2.6). We employed this algorithm to compute the drug response predictive performance of the joint signal estimated using the robust data alone (Figure 4A). Dividing the data in ten independent folds, we performed a cross-validation estimation as follows. For each train-test division of the data, we trained a Percolate instance on the 90% of the data, the training set containing GE and the robust data type. The resulting joint information was then used to train an ElasticNet model to predict drug response. The remaining 10% (test data) were then used to first estimate the joint signal, solely based on the robust data (Subsection 2.6). This joint signal was then used as input into the ElasticNet model to predict the response on this test set. Finally, we computed the predictive performance as indicated in Subsection 2.7.

**Figure 4:**
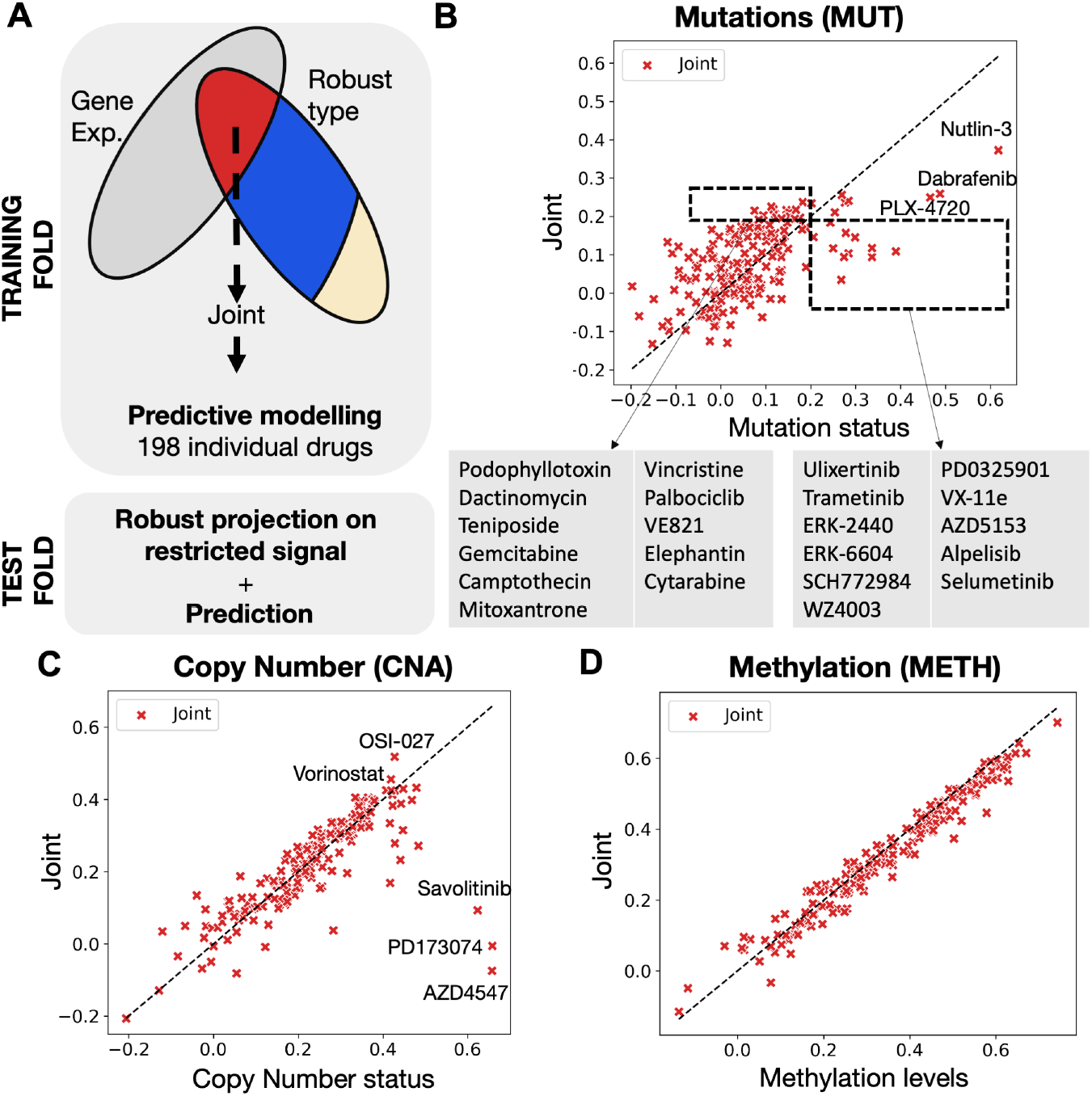
Robust-typebased signatures created from Percolate recapitulate drug response. (**A**) Schematic of the cross validation experiment. (**B**) Results for MUT with a special zoom on drugs predictive for joint but not robust (left) and for robust but not join (right). (**C**) Results for CNA. (**D**) Results for METH.

When analyzing results for MUT (Figure 4B), we first observe a clear drop in performance for the joint signal compared to the previous results (Figure 3B). This suggests that the GE portion of the joint signal (Equation 11) contains a significant portion of predictive signal, which is less well captured by our out-of-sample extension. Nonetheless, we observe that 11 drugs show a predictive performance above 0.2 for joint but not for the robust data. In contrast, 11 drugs show the opposite effect, including seven which target the MAPK pathway – MEK (Trametinib, PD0325901, Selumetinib) and ERK (ERK2440, ERK6604, Ulixertinib, SCH772984). BRAF inhibitors Dabrafenib and PLX-4720 also show a drop in performance. This suggests that constitutive activation of the MAPK pathway is not recapitulated by the joint signal. Nonetheless, the joint signal generated by Percolate helps increase performance for several poorly predictive drugs and is therefore of interest to study various response mechanisms. We then turned to CNA (Figure 4C) and observe a modest decrease in predictive performance compared to the performance on the original CNA profiles. Three drugs show a spectacular drop as the response can not be predicted by the joint signal – Savolitinib (cMET), PD173074 (FGFR) and AZD4547 (FGFR). In contrast, three drugs show improved performance for the joint signal – OSI-027 (mTOR), Navitoclax (HDAC) and Vincristine (tubulin). Finally, we repeated the experiment for METH (Figure 4D) and observe that predictive performances of the joint signal is remarkbly comparable to the predictive performance on the original METH data, with most drugs falling showing less than 2% relative performance difference (Supp. Figure 4C). Taken together, these results show that the joint signal recapitulates the drug response performance abilities of DNA-based measurements.

### 3.4 Study of genes contributing to the joint signals

We then set out to study the underlying mechanisms associated with the predictors derived from the robust data types (Subsection 3.3) which also lead to improved performance. For a given drug, we trained an ElasticNet model on the joint signal, yielding one regression coefficient per joint component. Using the relationship from Equation 11, we obtain a regression coefficient for each gene. A positive coefficient indicates that larger values of the saturated parameters, caused by a mutation or amplification of the supporting gene, are associated with resistance. In contrast, a negative coefficient indicates that larger values of the saturated parameters are associated with sensitivity.

For MUT, we studied the mode of action of three drugs for which the joint signal performs well (Figure 4B): Gemcitabine (Figure 5A), Vincristine (Figure 5B) and Palbociclib (Figure 5C). We observe that TP53 mutation status is associated with resistance to three drugs, concordant with earlier observations showing that TP53 mutant are more resistant to chemotherapy [14]. Resistance to Gemcitabine and Vincristine is also associated with KRAS and PI3KCA mutations, known for their proliferative potential [18, 10]. Interestingly, mutations in MYC and MAPK8IP2 are associated with sensitivity to these three drugs. Three other drugs show a drop in predictive performance on the joint signal as compared to the original signal: Nutlin-3, Dabrafenib and PLX-4720 (Figure 4B). We observe that the known targets of these drugs exhibit a large coefficient: TP53 for Nultin-3 (known resistance biomarker) and BRAF for Dabrafenib and PLX-4720 (Supp. Figure 5). These three drugs highlight a limitation of our approach: GLM-PCA generates scores which aggregates the contributions of several genes. Highly-specific drugs, like Nutlin-3 (Mdm2-inhibitor) or BRAF/MEK-inhibitors not only target a specific protein, but mutations in the target are excellent response predictors. Such cases do not benefit from the GLM-PCA aggregation as a single feature alone is predictive.

**Figure 5:**
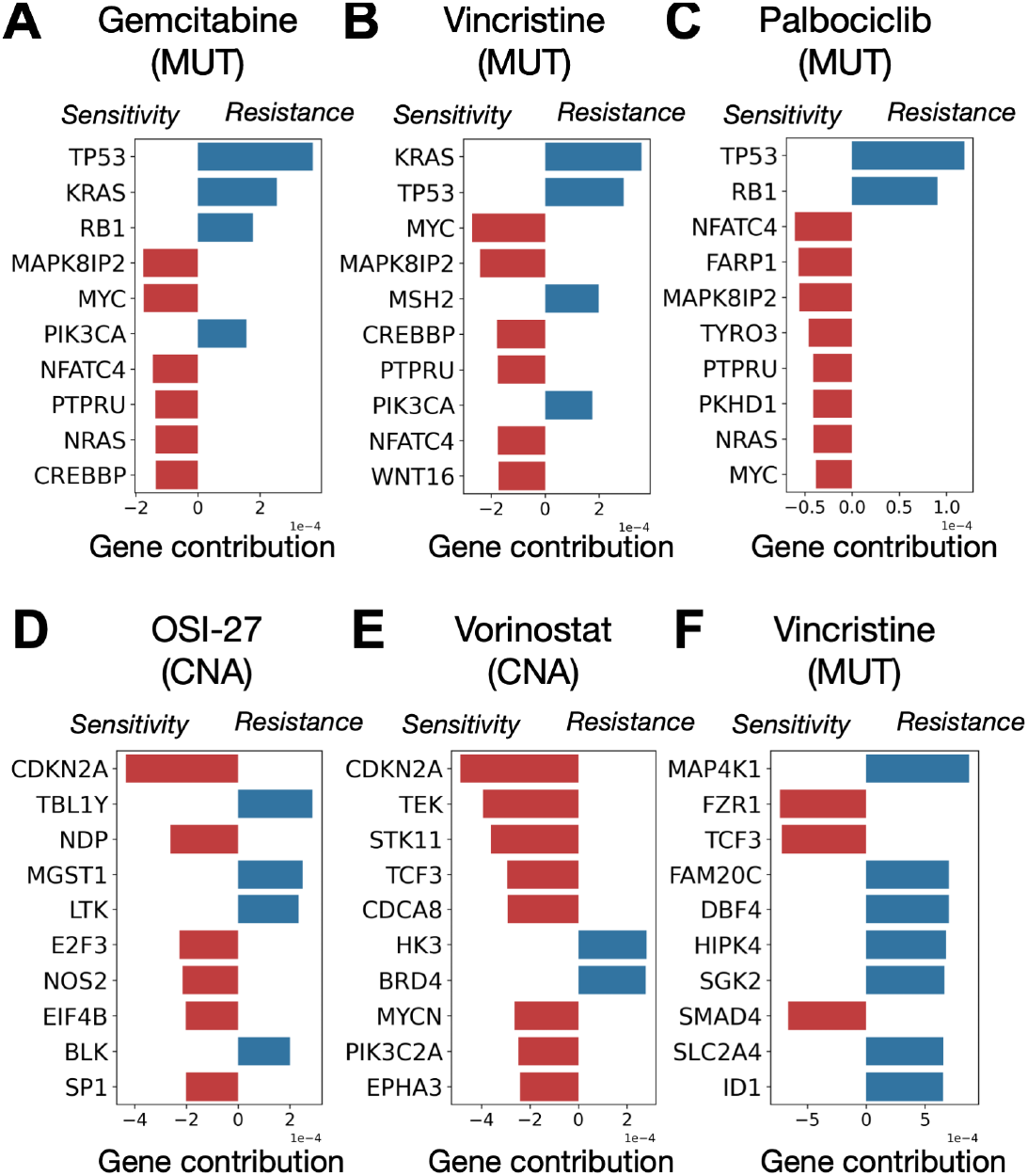
Study of joint signals contributing to improved performance. For each drug, we report the top 10 largest gene regression coefficients from the joint signal, in absolute values. We first analysed the joint biomarkers created from MUT data for Gemcitabine (**A**), Vincristine (**B**) and Palbociclib (**C**). We then turned to CNA-based signatures for OSI-27 (**D**), Vorinostat (**E**) and Vincristine (**F**).

Next we turned to CNA where three drugs: OSI-27 (Figure 5D), Vorinostat (Figure 5E) and Vincristine (Figure 5F), which all showed increased performance when the joint signal is employed as compared to the original CNA data. For both OSI-27 (mTORC1) and Vorinostat (HDAC), we observe that amplification of CDKN2A (p16) is associated with sensitivity. P16 acts as a tumor-suppressor by slowing down the early progression of the cell-cycle and its loss is here associated with resistance for these two drugs. Finally, Vincristine’s predictor shows that MAP4K1’s amplification as a predictor of resistance. Such result is coherent with what we observed for MUT (Figure 5B) where mutations on KRAS were associated with resistance.

### 3.5 Iterative application of Percolate deprives gene expression from predictive power

Finally, we questioned whether some signal predictive of drug response is still present in gene expression. To this end, we studied the GE signal after it has been stripped of all the signal it shares with MUT, METH or CNA. To remove all signal associated with robust data types from GE, we used Percolate iteratively on GE, starting with the *least* predictive data type (MUT), followed by CNA and ending with the *most* predictive data type (METH) (Figure 6A). Specifically, we first “percolate” GE through MUT to obtain an individual GE signal (not shared with MUT), which is then percolated through CNA to obtain a second GE individual signal, which is then finally percolated through METH, resulting in the individual GE signal we denote as *residual gene expression*. We finally assessed the predictive performance of this residual gene expression and compared it to the predictive performance of the original GE (Figure 6B, Subsection 2.7). We observe that no drug reaches a Pearson correlation above 0.16, indicative of a complete lack of predictive performance in the residual GE. This shows that removing the signal joint with DNA-based measurements deprives gene expression from any predictive ability.

**Figure 6:**
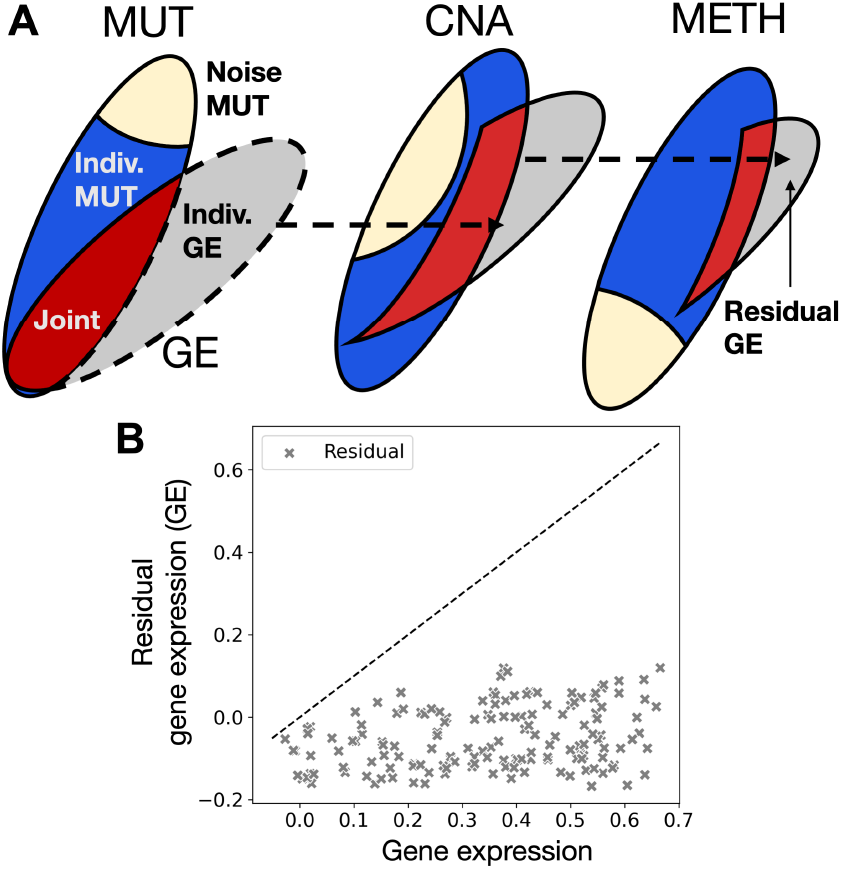
The signal joint with DNA-based measurements deprives gene expression from any predictive power. (**A**) Schematic of our iterative procedure to remove from GE any signal joint with robust data type. (**B**) Predictive performance of the resulting residual gene expression compared to the predictive performance of the complete gene expression.

## 4 Discussion

Designing multi-omics predictors of drug response has highlighted the existence of a trade-off between robust and predictive data types. To study this trade-off, we developed Percolate, a method which decomposes a pair of data types into a joint and an individual signal. After showing that the strength of the joint signal recapitulates the known topology between data types, we showed that the joint signal contains more predictive power than any robust data type alone. Exploiting our out-of-sample extension, we showed that the joint signal, computed from robust data types alone, recapitulates most of the predictive performance of each original robust signal. Finally, we showed that the gene expression signal predictive of drug response is fully captured by robust data types through Percolate.

Technically, Percolate extends JIVE in two different ways. First, by using GLM-PCA instead of PCA, we tailor the dimensionality reduction step to the specific data under consideration. Second, we developed an out-of-sample extension which allows to estimate the joint signal, even in the absence of one data-modality. For our analysis, we made use of standard distributions from the exponential family: Negative Binomial, Gamma, Beta or Bernoulli. Our implementation of GLM-PCA is versatile and any exponential family distribution can be employed in our framework, provided it can be auto-differentiated by PyTorch. Employing more complex distribution, like the inverse-gamma for copy-number is a fruitful avenue to improve on our methodology.

## Acknowledgements and Funding

We wish to thank Stavros Makrodimitris (TU Delft) for useful discussions, and the RHPC of the NKI for the computing infrastructure. This work was funded by ZonMw TOP grant COMPUTE CANCER (40-00812-98-16012).

## References

[1] N. Aben et al. TANDEM: A two-stage approach to maximize interpretability of drug response models based on multiple molecular data types. Bioinformatics, 32(17):i413–i420, 2016.

[2] N. Aben et al. ITOP: Inferring the topology of omics data. Bioinformatics, 34(17):i988–i996, 2018.

[3] H. Akaike. A new look at the statistical model identification. IEEE transactions on automatic control, 19(6):716–723, 1974.

[4] R. Argelaguet et al. Multi-Omics Factor Analysis—a framework for unsupervised integration of multi-omics data sets. Molecular Systems Biology, 14(6):1–13, 2018.

[5] M. Bersanelli et al. Methods for the integration of multi-omics data: Mathematical aspects. BMC Bioinformatics, 17(2), 2016.

[6] L. Cantini et al. Benchmarking joint multi-omics dimensionality reduction approaches for the study of cancer. Nature Communications, 12(1):1–12, 2021.

[7] M. Collins et al. A generalization of principal component analysis to the exponential family. NeurIPS, (1), 2002.

[8] J. C. Costello et al. A community effort to assess and improve drug sensitivity prediction algorithms. Nature Biotechnology, 32(12):1202–1212, 2014.

[9] J. M. Dempster et al. Gene expression has more power for predicting in vitro cancer cell vulnerabilities than genomics. bioRxiv, 1:1–42, 2020.

[10] E. A. Eklund et al. Kras mutations impact clinical outcome in metastatic non-small cell lung cancer department of surgery. 2022.

[11] Q. Feng et al. Angle-based joint and individual variation explained. Journal of Multivariate Analysis, 166:241–265, 2018.

[12] M. Ghandi et al. Next-generation characterization of the Cancer Cell Line Encyclopedia. Nature, 569(7757):503–508, 2019.

[13] D. Hanahan et al. Hallmarks of cancer: The next generation. Cell, 144(5):646–674, 2011.

[14] K. Hientz et al. The role of p53 in cancer drug resistance and targeted chemotherapy. Oncotarget, 8(5):8921–8946, 2017.

[15] M. Hoogstraat et al. Genomic and transcriptomic plasticity in treatment-naïve ovarian cancer. Genome Research, 24(2):200–211, 2014.

[16] F. Iorio et al. A Landscape of Pharmacogenomic Interactions in Cancer. Cell, 2016.

[17] I. S. Jang et al. Systematic assessment of analytical methods for drug sensitivity prediction from cancer cell line data. In Pacific Symposium for Biocomputing, volume 23, pages 1–7, 2013.

[18] S. T. Kim et al. Impact of KRAS mutations on clinical outcomes in pancreatic cancer patients treated with first-line gemcitabine-based chemotherapy. Molecular Cancer Therapeutics, 10(10):1993–1999, 2011.

[19] Y. Kim et al. WON-PARAFAC: a genomic data integration method to identify interpretable factors for predicting drug-sensitivity in-vivo. pages 1–30.

[20] V. N. Kristensen et al. Kristensen - Principles and methods of integrative genomic analyses in cancer.pdf. Nature Reviews Cancer, 14:299–313, 2014.

[21] A. J. Landgraf et al. Dimensionality reduction for binary data through the projection of natural parameters. Journal of Multivariate Analysis, 180(1999), 2020.

[22] N. Lawrence. Probabilistic non-linear principal component analysis with Gaussian process latent variable models. Journal of Machine Learning Research, 6:1783–1816, 2005.

[23] J. Li et al. Simple exponential family PCA. IEEE Transactions on Neural Networks and Learning Systems, 24(3):485–497, 2013.

[24] Y. Li et al. A review on machine learning principles for multi-view biological data integration. Briefings in bioinformatics, 19(2):325–340, 2018.

[25] L. T. Liu et al. ePCA: High dimensional exponential family PCA. Annals of Applied Statistics, 12(4):2121–2150, 2018.

[26] E. F. Lock et al. Joint and individual variation explained (JIVE) for integrated analysis of multiple data types. Annals of Applied Statistics, 7(1):523–542, 2013.

[27] I. Martincorena et al. Somatic mutation in cancer and normal cells. Science, 349(6255):1483–1489, 2015.

[28] H. L. McLeod. Cancer pharmacogenomics: Early promise, but concerted effort needed. Science, 340(6127):1563–1566, 2013.

[29] M. Meghwanshi et al. McTorch, a manifold optimization library for deep learning. pages 1–5, 2018.

[30] A. Paske et al. Automatic differentiation in prose. In NeurIPS’, 2017.

[31] E. J. Pitman. Sufficient Statistics and Intrinsic Accuracy. Mathematical Proceedings of the Cambridge Philosophical Society, 32(4):567–579, 1936.

[32] M. G. Rees et al. Correlating chemical sensitivity and basal gene expression reveals mechanism of action. Nature Chemical Biology, 12(2):109–116, 2016.

[33] M. V. Relling et al. Pharmacogenomics in the clinic. Nature, 526(7573):343–350, 2015.

[34] M. D. Robinson et al. edgeR: A Bioconductor package for differential expression analysis of digital gene expression data. Bioinformatics, 26(1):139–140, 2009.

[35] C. Sagonas et al. Robust joint and individual variance explained. CVPR, 2017:5739–5748, 2017.

[36] H. Sharifi-Noghabi et al. MOLI: Multi-omics late integration with deep neural networks for drug response prediction. Bioinformatics, 35(14):i501–i509, 2019.

[37] H. Shu et al. D-CCA: A Decomposition-Based Canonical Correlation Analysis for High-Dimensional Datasets. Journal of the American Statistical Association, 115(529):292–306, 2020.

[38] A. M. Smith et al. Standard machine learning approaches outperform deep representation learning on phenotype prediction from transcriptomics data. BMC Bioinformatics, 21(1):1–18, 2020.

[39] M. E. Tipping et al. Probabilistic Principal Component Analysis. Journal of the Royal Statistical Society, Series B, 61(3):611–622, 1999.

[40] F. W. Townes, S. C. Hicks, M. J. Aryee, and R. A. Irizarry. Feature selection and dimension reduction for single-cell RNA-Seq based on a multinomial model. Genome Biology, 20(1):1–16, 2019.

[41] B. Wang et al. Similarity network fusion for aggregating data types on a genomic scale. Nature Methods, 11(3):333–337, 2014.

[42] H. Zou et al. Regularization and variable selection via the elastic net Hui. Journal of the Statistical Society, Series B, 67(2):301–320, 2005.

