## Supplementary Material for "Percolate: an exponential family JIVE model to design DNA-based predictors of drug response"

#### 1 Negative binomial parametrization

##### 1.1 Common parametrization

The Negative Binomial distribution is commonly defined using two parameters:

- $r > 0$, called the **inverse-dispersion**.
- $p \in [0, 1]$, called the **success probability**.

The negative binomial distribution models the number of successes observed when consecutive Bernoulli experiments with probability $p$ are carried out and stopped after $r$ failures. Its probability distribution is defined on $\mathbb{N}$ as follows,

$$\forall k \in \mathbb{N}, \quad \mathbb{P}(X = k) \;=\; \frac{\Gamma(k+r)}{\Gamma(k+1)\Gamma(r)} (1-p)^r p^k, \tag{1}$$

where $\Gamma$ stands for the Gamma-distribution defined as

$$\forall z > 0, \quad \Gamma(z) \;=\; \int_0^{+\infty} t^{z-1} e^{-t} dt. \tag{2}$$

If we re-write Equation (1) in an exponential form, we obtain:

$$\forall k \in \mathbb{N}, \quad \mathbb{P}(X = k) \;=\; \frac{\Gamma(k+r)}{\Gamma(k+1)\Gamma(r)} \exp\left[r \ln(1-p) + k \ln p\right]. \tag{3}$$

In Equation (3), we easily recognise an exponential form when $r$ is fixed with $p$ as parameters. However, this formulation is computationally challenging, as observed values of $p$ tends to accumulate around 1. This leads to instabilities in the optimisation and motivate the utilisation of another parametrization.

##### 1.2 Parametrization employed in our approach

We turned to the parametrization proposed by *Risso et al* [3], which construction we present here. This parametrization stems from the observation that, if $Z$ follows a Negative Binomial distribution with parameters $p$ and $r$, then $\mathbb{E}(Z) = \frac{pr}{1-p}$. This expectation belongs to $\mathbb{R}^+$ which still contains one constraint. This constraint can easily be alleviated by log-transform, leading to the following parametrization:

$$\theta \;=\; \ln \frac{pr}{1-p}. \tag{4}$$

Equivalently, we have $p = \frac{r}{e^\theta + r}$, which combined with Equation (3) yields the exponential form presented in the main text.

##### 1.3 Technical implementation

The Negative Binomial distribution $\text{NB}(\theta, r)$ depends on two parameters $\theta \in \mathbb{R}$ and $r > 0$. When the $r$ parameter, called **inverse-dispersion**, is fixed, the Negative Binomial distribution belongs to the exponential family. We use the parametrization used by *Risso et al* [3] which yields, with fixed-parameter $r > 0$, the exponential-family functions explicited in Supp. Table 1. We compute for each gene (feature) $j \in \{1, .., p\}$ a dispersion parameter $r_j$ using DESeq2 [2]. Once these dispersion parameters set, the parametrization can be exploited in Percolate.

#### 2 Beta parametrization

##### 2.1 Common parametrization

The Beta distribution is commonly parametrized by two parameters, $\alpha > 0$ and $\beta > 0$, called **shape** parameters. If a random-variable $Z$ follows the Beta distribution with parameters $\alpha$ and $\beta$, then its probability density function $f(\cdot; \alpha, \beta)$ is defined as

$$\forall z \in [0,1], \quad f(z; \alpha, \beta) \;=\; \frac{\Gamma(\alpha+\beta)}{\Gamma(\alpha)\Gamma(\beta)} z^{\alpha-1}(1-z)^{\beta-1}. \tag{5}$$

Noting that $\mathbb{E}[Z] = \frac{\alpha}{\alpha+\beta}$, we can use another parametrization which brings the saturated parameters to be the data-expectation, as is the case for other distributions like Gaussian, Bernoulli, Poisson or Negative-Binomial.

##### 2.2 Parametrization employed in our approach

Using the parametrization from Equation (5), we define the two parameters $\theta$ and $\nu$ as:

$$\theta \;= \frac{\alpha}{\alpha+\beta} \quad \text{and} \quad \nu \;= \alpha+\beta. \tag{6}$$

The re-parametrization from Equation (6) can be reversed, yielding $\alpha = \theta\nu$ and $\beta = (1-\theta)\nu$. The probability density function from Equation (5) then becomes

$$\forall z \in [0,1], \quad f(z; \theta, \nu) \;=\; \frac{1}{1-z} \exp\left[\begin{bmatrix} \theta\nu \\ (1-\theta)\nu \end{bmatrix}^T \begin{bmatrix} \ln z \\ \ln(1-z) \end{bmatrix} - \ln \frac{\Gamma(\theta\nu)\Gamma((1-\theta)\nu)}{\Gamma(\nu)}\right]. \tag{7}$$

From Equation (7), one can easily derive the $A$, $T$ and $\eta$ functions listed in Table 1 of the main manuscript.

##### 2.3 Computing saturated parameters

Let $x \in [0,1]$ and $\nu > 0$, the saturated parameter $\widetilde{\theta}$ solves the following equation

$$\frac{df}{d\theta}\left(x; \widetilde{\theta}, \nu\right) \;=\; 0. \tag{8}$$

We denote by $\psi$ the **digamma** function defined as the derivative of the log-Gamma function:

$$\forall t > 0, \quad \psi(t) \quad = \quad \frac{d \ln \Gamma}{dt}(t) \quad = \quad \frac{1}{\Gamma(t)} \frac{d\Gamma}{dt}(t). \tag{9}$$

Combining Equations (7) and (9), and noting that $r > 0$, the saturated parameter $\widetilde{\theta}$ is the solution of

$$\ln \frac{x}{1-x} \;+\; \psi((1-\theta)\nu) \;-\; \psi(\theta\nu) \quad = \quad 0. \tag{10}$$

We define $g$ as $g : \theta \mapsto \ln \frac{x}{1-x} \;+\; \psi((1-\theta)\nu) \;-\; \psi(\theta\nu)$. We have:

$$\forall \theta > 0, \quad \frac{dg}{d\theta}(\theta) \quad = \quad -\nu\left[\psi^{(1)}((1-\theta)\nu) \;+\; \psi^{(1)}(\nu\theta)\right], \tag{11}$$

where $\psi^{(1)}$ represent the first-order derivative of the digamma function. $\psi^{(1)}$ can be efficiently computed using its integral representation:

$$\forall z > 0, \quad \psi^{(1)}(z) \; = \; -\int_0^1 \frac{t^{z-1}}{1-t} \ln t \; dt \tag{12}$$

Using Equation (12), we obtain

$$\forall \theta > 0, \quad \frac{dg}{d\theta}(\theta) \quad = \quad \nu \left[ \int_0^1 \frac{t^{\theta\nu - 1} + t^{\nu - \theta\nu - 1}}{1-t} \ln t \; dt \right] \quad \geq \quad 0. \tag{13}$$

Using the monotonicity of $g$, we easily solve Equation (8) by means of a dichotomy. A Newton method would have also been possible, but would have required to set a learning rate. Our approach allows to avoid such additional hyper-parameters.

##### 2.4 Technical implementation

The Beta distribution $B(\theta, \nu)$ depends on two parameters $\theta \in [0,1]$ and $\nu > 0$ and belongs to the exponential family, defined, for $x \in [0,1]$ as in Supp. Table 1. Since the Beta distribution has two natural parameters, we took inspiration from the Negative Binomial and fixed $\nu$ at the gene-level: since $\theta$ can be intuitively understood as the mean of the distribution, we reasoned that it would act as a good parameter for GLM-PCA. We computed the parameters $\nu_1, .., \nu_p$ by maximizing the likelihood for each gene and used the resulting distributions in Percolate.

#### 3 Bernoulli parametrization

The natural parameters from the Bernoulli distribution are either $-\infty$ or $\infty$ and thus not computationally tractable. Following *Landgraf et al* [1], we employ a thresholding at 25 for all entries of the matrix $\widetilde{\theta}$ ; this choice did not impact results (Supp. Figure 2).

#### 4 Proof of Proposition 2.4

Using the differentiability of $A$ and $\eta$, we obtain, for $\Theta \in \mathbb{R}^{n \times p \times q}$ and $i \in \{1, ..n\}$ and $j \in \{1, ..p\}$

$$(\nabla L(\Theta))_{i,j,\cdot} \; = \; \nabla A(\Theta_{i,j,\cdot}) \; - \; J_\eta (\Theta_{i,j,\cdot})^T \, T(X_{i,j}). \tag{14}$$

The minimum of the negative log-likelihood is obtained when the differential is zero, which is obtained when

$$\nabla A(\Theta_{i,j,\cdot}) \; = \; J_\eta (\Theta_{i,j,\cdot})^T \, T(X_{i,j}). \tag{15}$$

Using the invertibility of the derivatives and the vector structure of $\nabla A(\Theta_{i,j,\cdot})$, we obtain

$$\nabla A(\Theta_{i,j,\cdot}) \, J_\eta (\Theta_{i,j,\cdot})^{-1} \; = \; T(X_{i,j}) \tag{16}$$

By invertibility of $\eta$ and using chain rule, we obtain

$$J_{A \circ \eta^{-1}} (\eta(\Theta_{i,j,\cdot})) \; = \; T(X_{i,j}), \tag{17}$$

which then leads to

$$\Theta_{i,j\cdot} \; = \; \eta^{-1} \left[ \left( J_{A \circ \eta^{-1}} \right)^{-1} (T(X_{i,j})) \right] \; = \; \eta^{-1} \circ \left( J_{A \circ \eta^{-1}} \right)^{-1} \circ T(X_{i,j}). \tag{18}$$

#### 5 Proof of Theorem 2.7

**Lemma 5.1.** *Let $V \in \mathbb{R}^{p \times d}$ such that $V^T V = I_d$, $\Sigma \in \mathbb{R}^{d \times d}$ be a diagonal matrix and $\mu \in \mathbb{R}^p$, then*

$$\min_{\substack{U \in \mathbb{R}^{n \times d}, \Sigma \in \mathbb{R}^{d \times d} \\ U^T U = I_d}} \mathcal{L}\left( U \Sigma V^T + 1_n \mu^T \; ; \; X, \mathcal{E} \right) \quad \geq \quad \mathcal{L}\left( \left( \widetilde{\Theta}(X; \mathcal{E}) - 1_n \mu^T \right) V V^T + 1_n \mu^T \; ; \; X, \mathcal{E} \right) \tag{19}$$

*Proof.* Observe that for $U \in \mathbb{R}^{n\times d}$ such that $U^T U = I_d$ and $\Sigma \in \mathbb{R}^{d\times d}$ diagonal matrix, $U\Sigma \in \mathbb{R}^{p\times d}$. This directly entails the following inequality:

$$\min_{\substack{U\in\mathbb{R}^{n\times d},\Sigma\in\mathbb{R}^{d\times d}\\ U^TU=I_d}} \mathcal{L}\left(U\Sigma V^T + 1_n\mu^T \; ; \; X,\mathcal{E}\right) \quad \geq \quad \min_{U\in\mathbb{R}^{n\times d}} \mathcal{L}\left(UV^T + 1_n\mu^T \; ; \; X,\mathcal{E}\right). \tag{20}$$

Deriving the functions defined at the right-hand side of Equation 20 and equating the first derivative to zero, the matrix $U^*$ minimizing the right-hand-side problem satisfies

$$V^T\left[\nabla A\left(U^*V^T + 1_n\mu^T\right) \; - \; J_\eta\left(U^*V^T + 1_n\mu^T\right)^T T(X)\right] \; = \; 0, \tag{21}$$

using a matrix element-wise notation. Noting that $V \neq 0$ and using the same computation as in the proof of Proposition 2.4, we obtain:

$$U^*V^T + 1_n\mu^T \quad = \quad \eta^{-1}\left[\left(J_{A\circ\eta^{-1}}\right)^{-1}(T(X))\right]. \tag{22}$$

Combining Equation 22 with Proposition 2.4, we obtain

$$U^*V^T \quad = \quad \widetilde{\Theta}(X;\mathcal{E}) \; - \; 1_n\mu^T. \tag{23}$$

Finally, multiplying left- and right-hand sides by $V$ and exploiting that $V^TV = I_d$, we obtain:

$$U^* \quad = \quad \left(\widetilde{\Theta}(X;\mathcal{E}) \; - \; 1_n\mu^T\right)V. \tag{24}$$

Combining Equations 20 and 24 yields the result. ■

Theorem 2.7 follows directly by application of Supp. Lemma 5.1.

### 6 Proof of Theorem 2.11

Without loss of generality, we restrict to non-singular directions, i.e. $\Sigma_{i,i} > 0 \forall i$. Using the decomposition $V_M = \left[V_{M,A}^T \; V_{M,B}^T\right]^T$ and the fact that $V_M^T V_M = I$ (by construction), we obtain directly:

$$\begin{aligned} U_M \quad &= \quad \left[\widetilde{U}_A, \widetilde{U}_B\right]\begin{bmatrix} V_{M,A} \\ V_{M,B}\end{bmatrix}\Sigma_M^{-1} \\ &= \quad \widetilde{U}_A V_{M,A}\Sigma_M^{-1} \; + \; \widetilde{U}_B V_{M,B}\Sigma_M^{-1}. \end{aligned} \tag{25}$$

If we further note that $\left(\widetilde{\theta}_A - \widetilde{\mu}_A\right)\widetilde{V}_A^T\widetilde{V}_A = \widetilde{U}_A\Sigma_A W_A^T$ and $\left(\widetilde{\theta}_B - \widetilde{\mu}_B\right)\widetilde{V}_B^T\widetilde{V}_B = \widetilde{U}_B\Sigma_B W_B^T$ (Definition 2.8), we obtain:

$$\begin{aligned} \widetilde{U}_A \quad &= \quad \left(\widetilde{\theta}_A - \widetilde{\mu}_A\right)\widetilde{V}_A^T\widetilde{V}_A W_A\Sigma_A^{-1} \\ \widetilde{U}_B \quad &= \quad \left(\widetilde{\theta}_B - \widetilde{\mu}_B\right)\widetilde{V}_B^T\widetilde{V}_B W_B\Sigma_B^{-1} \end{aligned}. \tag{26}$$

Combining Equations (25) and (26) yields the desired formula.

| Name | $\mathcal{X}$ | $T(x)$ | $\eta(\theta)$ | $A(\theta)$ | $g^{-1}$ | $\mathbb{E}\left[T\left(X\right)\right]$ |
|---|---|---|---|---|---|---|
| Gaussian | $\mathbb{R}$ | $x$ | $\theta$ | $\theta^2/2$ | $x$ | $\theta$ |
| Bernoulli | $\{0,1\}$ | $x$ | $\theta$ | $\log\left(1+e^{\theta}\right)$ | $+\infty$ if $x=1$ else $-\infty$ | $\left(1+e^{-\theta}\right)^{-1}$ |
| Poisson | $\mathbb{R}_+$ | $x$ | $\theta$ | $e^{\theta}$ | $\log\left(x\right)$ | |
| Negative Binomial | $\mathbb{R}_+$ | $x$ | $\log\frac{r}{e^{\theta}+r}$ | $r\log\left(1+re^{-\theta}\right)$ | $2\log r-\log x$ | $r^2e^{-\theta}$ |
| Beta | $[0,1]$ | $\begin{bmatrix}\log x\\ \log\left(1-x\right)\end{bmatrix}$ | $\begin{bmatrix}\theta\nu\\ \left(1-\theta\right)\nu\end{bmatrix}$ | $\log\frac{\Gamma(\theta\nu)\Gamma((1-\theta)\nu)}{\Gamma(\nu)}$ | untractable | $\begin{bmatrix}\psi\left(\theta\nu\right)-\psi\left(\nu\right)\\ \psi\left(\left(1-\theta\right)\nu\right)-\psi\left(\nu\right)\end{bmatrix}$ |
| Log-normal | $\mathbb{R}_+$ | $\begin{bmatrix}\log x\\ \left(\log x\right)^2\end{bmatrix}$ | $\begin{bmatrix}\theta/\nu^2\\ -1/\left(2\nu^2\right)\end{bmatrix}$ | $\frac{\theta^2}{2\nu^2}+\log\nu$ | $\log x$ | $\begin{bmatrix}\theta\\ \theta^2-\sigma^2\end{bmatrix}$ |
| Gamma | $\mathbb{R}_+$ | $\begin{bmatrix}\log x\\ x\end{bmatrix}$ | $\begin{bmatrix}\theta\\ -\nu\end{bmatrix}$ | $\log\Gamma\left(\theta+1\right)-\left(\theta+1\right)\log\nu$ | $\psi^{-1}\left(\log\left(\nu x\right)\right)-1$ | $\begin{bmatrix}\psi\left(\theta+1\right)-\log\nu\\ -\left(\theta+1\right)/\nu\end{bmatrix}$ |

Table 1: **Exponential family distributions**. Gaussian distribution is assumed to have unit variance. The inverse-dispersion parameter $r$ is fixed for the Negative Binomial. For the Beta and the log-normal distributions, $\nu$ refers to the second parameter computed at the feature-level.

### 7 Supplementary figures

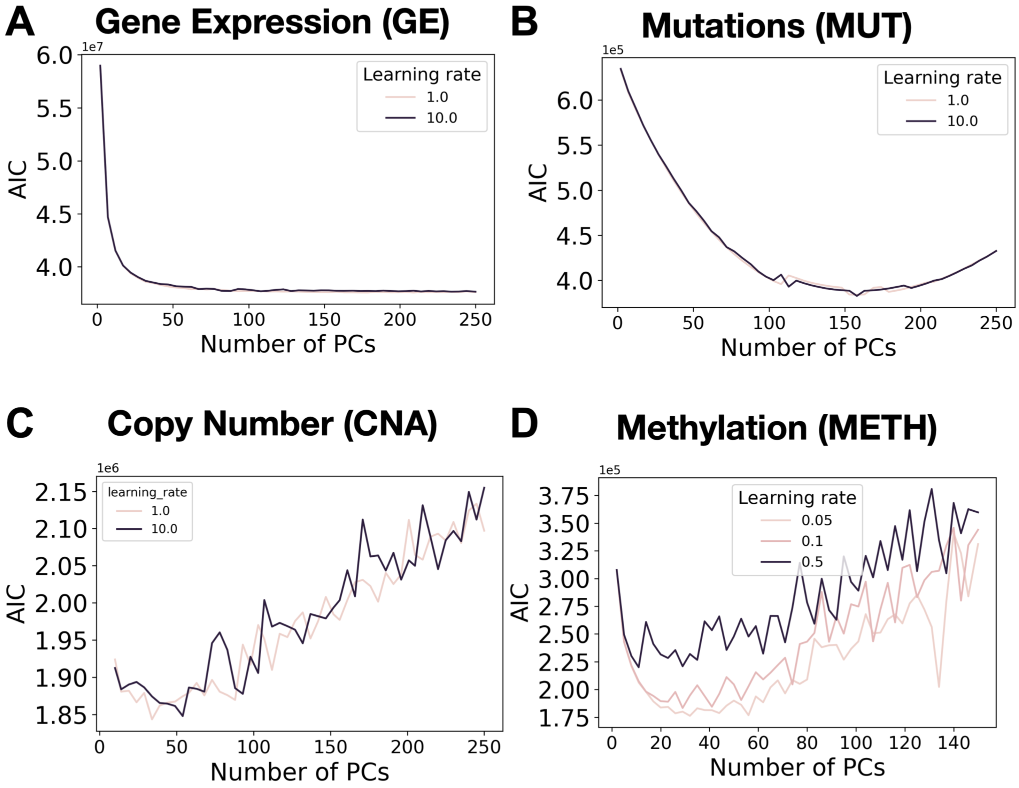


Figure 1: **Model selection for each data-type.** For each data-type we compute the AIC on the complete dataset for various hyper-parameters and report the best performance (i.e., lowest AIC) per number of components. This model selection wass performed for (**A**) gene expression (negative binomial), (**B**) mutation (bernoulli), (**C**) copy number (gamma) and (**D**) methylation (beta).

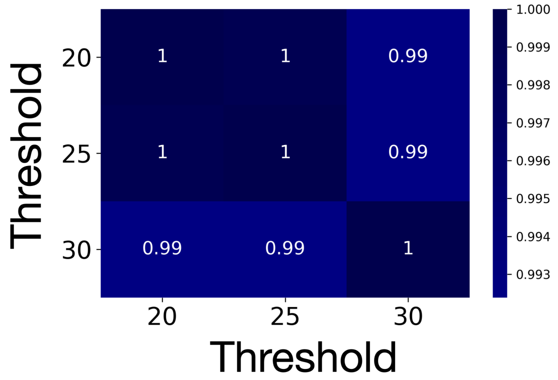


Figure 2: **Impact of threhsolding for mutations.** The infinity values in mutations are replaced by a large number, called **threshold**. To measure the impact of this parameter on Bernoulli-GLM-PCA, we used the optimal number of PCs set in Supp. Figure 1 and fit a GLM-PCA for different threshold values. We then compared the loadings (Equation 4) between experiments by reporting the average singular value of the dot-product matrix ; i.e. if $V_{20} \in \mathbb{R}^{d \times p}$ and $V_{30} \in \mathbb{R}^{d \times p}$ contain the loadings for threshold=20 and threshold=30 respectively, we compute the spectrum (singular values) of $V_{20}V_{30}^T$ and returns the mean. Values can range from 0 to 1. For threshold 40 or above, GLM-PCA does not converge due to gradient explosion.

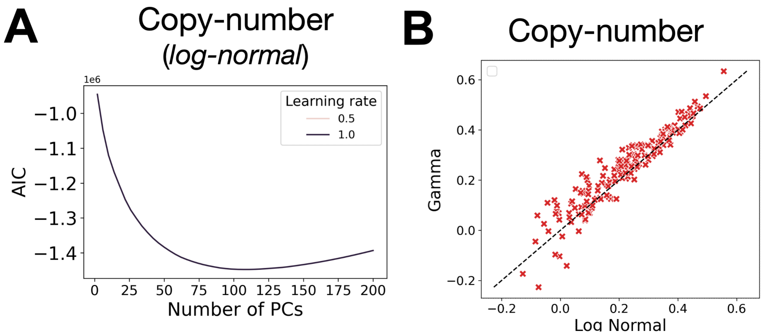


Figure 3: **Copy-number results with log normal distribution.** (**A**) AIC model selection for copy-number with log normal noise model (**B**) Comparison of predictive performance for the joint signal between CNA and GE for log-normal (x-axis) and gamma (y-axis) distribution. Each dot represents a single drug.

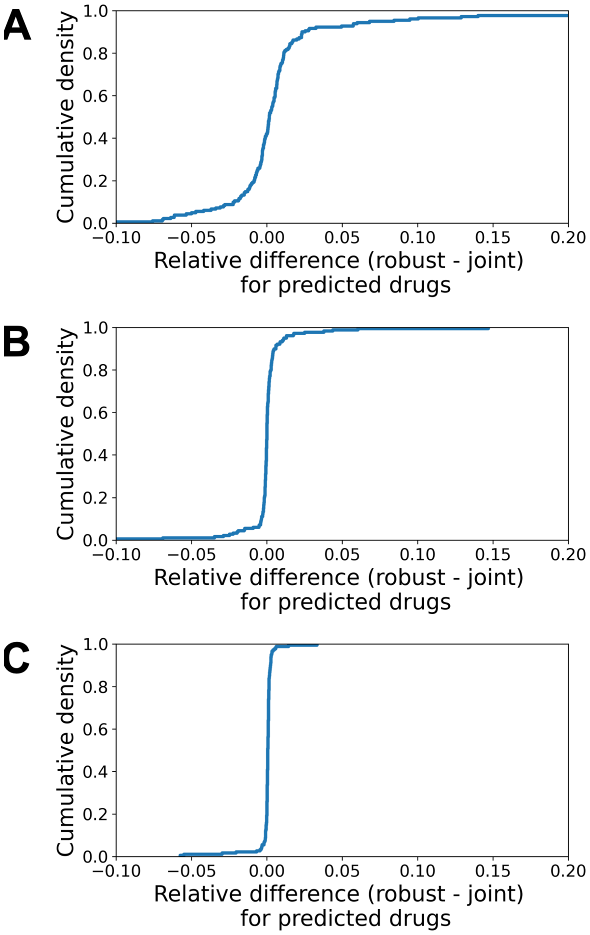


Figure 4: **Relative difference in predictive performance between joint signal (full cross-validation) and robust cell-view.** For each data-type, we computed the relative difference in predictive performance between the joint ($p_j$) and cell-view ($p_r$) as $(p_j - p_r)/p_j$. We report results for (**A**) mutations, (**B**), copy-number and (**C**) methylation.

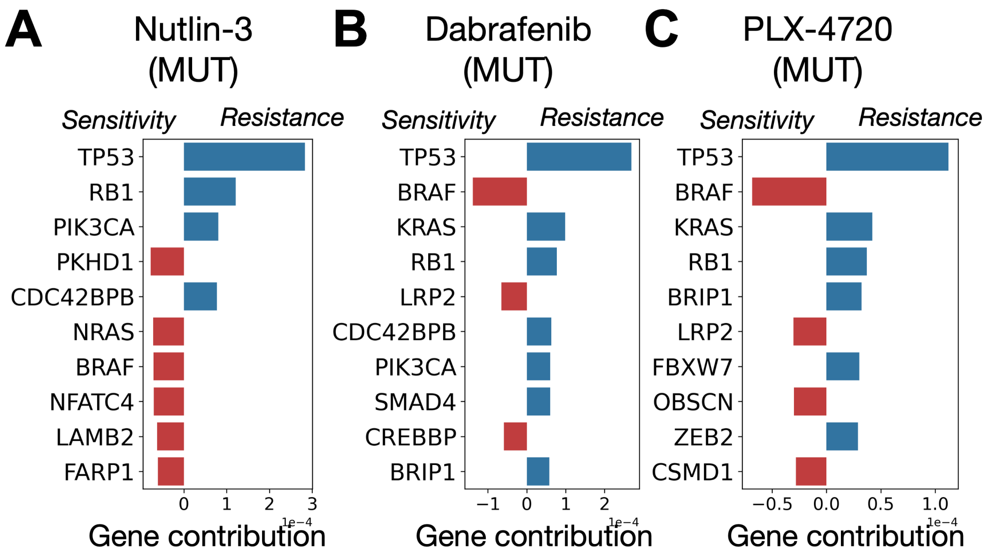


Figure 5: **Analysis of mutation joint-signal predictors.** Supporting Figure 5, we here report the predictors build on the joint signal between mutation and gene expression (solely based on mutations). (**A**) Nutlin-3, (**B**) Dabrafenib and (**C**) PLX-4720.

### References

[1] Andrew J. Landgraf and Yoonkyung Lee. Dimensionality reduction for binary data through the projection of natural parameters. *Journal of Multivariate Analysis*, 180(1999), 2020.

[2] Michael I. Love, Wolfgang Huber, and Simon Anders. Moderated estimation of fold change and dispersion for RNA-seq data with DESeq2. *Genome Biology*, 15(12):1–21, 2014.

[3] Davide Risso, Fanny Perraudeau, Svetlana Gribkova, Sandrine Dudoit, and Jean Philippe Vert. A general and flexible method for signal extraction from single-cell RNA-seq data. *Nature Communications*, 9(1):1–17, 2018.
